# Morphological structure of ant assemblages in tropical and temperate forests

**DOI:** 10.1101/065417

**Authors:** Rogério R. Silva, Israel Del Toro, Carlos Roberto F. Brandão, Aaron M. Ellison

## Abstract

Morphological variation in co-occurring species often is used to infer species assembly rules and other processes structuring ecological assemblages. We compared the morphological structure of ant assemblages in two biogeographic regions along two extensive latitudinal gradients to examine common patterns and unique characteristics of trait distribution. We sampled ant assemblages along extensive latitudinal gradients in Tropical Atlantic Forest in eastern Brazil and temperate forests in the eastern United States. We quantified 14 morphological traits related to the ecology and life history of each of 599 ant species and defined the morphological space occupied by different ant assemblages. Null models were used to test whether tropical and temperate ant assemblages differed from random expectation in morphological structure. Correlations between traits and climate were used to infer associations between habitat characteristics and morphological space occupied by ant assemblages. Tropical ant assemblages had higher morphological diversity and variation in the space of occupied morphospace, whereas temperate assemblages had higher variance in size. Although tropical ant assemblages had smaller morphological distances among species, species packing (i.e., mean nearest-neighbor distance) did not differ between regions. Null model analysis revealed scant evidence of habitat filtering or niche differentiation within assemblages. Different traits had different means, variances, skewness, and kurtosis values along each environmental gradient. Mean trait values within assemblages were associated mainly with region and correlated with temperature but trait variances had more complex responses to climate, including interactions between temperature and precipitation in the models. The higher functional diversity in tropical ant assemblages occurs by expansion of the morphospace rather than through an increase in species packing. Different traits vary independently along environmental gradients. Analysis of individual traits together with categorization of the moments of trait distributions (statistical central tendencies) provide new directions for quantifying morphological diversity in ant assemblages.

## Introduction

Contemporary community ecology is strongly focused on functional traits, what constrains them, and their relationship with environmental variables and community structure (Cadotte et al. 2013, Winemiller et al. 2015). An increasing number of studies suggest that examining functional traits of individual organisms may help to construct a mechanistic and predictive framework for the study of species distribution and co-occurrence (Cadotte et al. 2013). Trait diversity, which is closely related to functional diversity, increasingly has been used as a measure of phenotypic diversity in species assemblages (McGill et al. 2015). However, few studies have evaluated how morphological diversity is distributed along extensive latitudinal or environmental gradients, particularly in highly diverse invertebrate assemblages (Arnan et al. 2014, Yates et al. 2014, Silva and Brandão 2014). Without such analyses, it remains difficult to identify or test mechanistic hypotheses regarding the geographic distribution of biodiversity in light of the distribution of organismal functions (Swenson et al. 2012). A better understanding of the relationships between traits and geographic or environmental variables also is critical for improving predictions of functional and composition responses to ongoing global climatic change (Fortunel et al. 2014).

The morphospace of an assemblage can be defined as the distribution of morphological trait values within it (Ricklefs and Travis 1980). Quantifying the volume, overlap, and packing of morphological trait space generally improves our understanding of how different ecological processes structure morphological diversity and ecological strategies (Mouillot et al. 2005, Lamanna et al. 2014). For example, studies of morphological variation have revealed several interesting patterns in the occupation of niche space (Stevens et al. 2006, Inward et al. 2011, Ricklefs, 2012), morphological disparity between assemblages (how the variety of body plans within a taxon fills a morphospace) (McClain et al. 2004), responses of assemblage structure to habitat complexity (Willis et al. 2005, Montaña and Winemiller, 2010), and convergent or divergent patterns of morphology among continents (Ricklefs and Travis 1980, Winemiller 1991, Inward et al. 2011, Ricklefs 2012, Montaña et al. 2014). Morphological data also have been used to test the relative importance of environmental filtering and limiting similarity in structuring species assemblages (e.g., Cadotte et al. 2013, 2015).

The trait approach may be particularly useful for interpreting patterns of diversity of taxa such as arthropods that are hyperdiverse, relatively small in size, and poorly studied. For many arthropod groups, morphology of functionally important traits is the most easily obtainable, repeatable, and accurately measurable dimension associated with ecological diversity (Pie and Traniello 2007, Yates et al. 2014). Among the arthropods, ants are an ideal taxonomic group for investigating relationships between morphology, traits, and assemblage structure because they (i) are abundant, comprising the dominant fraction of animal biomass in most terrestrial ecosystems (Wilson and Hölldobler, 2005), (ii) perform a range of important ecosystems functions (Del Toro et al. 2012), (iii) exhibit a very large morphological diversity at very small scales (Silva and Brandão 2010), and (iv) vary in composition along environmental gradients (Gotelli and Ellison 2002). In addition, many studies have shown that ant species have specific traits that are correlated with environmental conditions (Wiescher et al. 2012), macro- or microhabitat structure (Kaspari and Weiser 1999, Farji-Brener et al. 2004, Gibb and Parr 2010, Yates et al. 2014) and resource exploitation (Retana et al. 2015). Finally, recent research with stable isotopes has identified evidence of a relationship between ant morphology and their trophic position within food webs (Gibb and Cunningham 2013, Gibb et al. 2015b).

Here, we present the first broad-scale description of the morphological differentiation of functional traits between temperate and tropical ant assemblages. We used three approaches to quantify morphological differentiation. First, we used diversity metrics to estimate the amount of morphological space occupied by temperate and tropical ant assemblages. Second, we combined multivariate statistics and null model analyses to ask whether morphological differentiation within assemblages was greater than expected, given their regional species pools. Third, we contrasted the moments of trait distributions between the two regions.

We combined these analytical approaches to test the following predictions: 1) Because of higher species diversity in the tropics and many more specialized predators, tropical and temperate-zone ant assemblages differ in morphological diversity and structure; 2) Because temperate forests experience harsher and more variable climates throughout the year, temperate ant assemblages should have a stronger imprint of habitat filtering (morphological clustering); and 3) Ecological traits of ant assemblages are correlated with climate. Our data base of functional traits is one of the largest compiled to date for ants and includes a large number of species-specific morphological traits from different assemblages across two extensive environmental gradients. We identified both similarities and differences in morphological structure of ant assemblages in tropical and temperate ant faunas and illustrate that analysis of individual traits can reveal unexpected patterns in relationships between traits, environmental gradients, and changes in land-use.

## Material and Methods

### Data Analysis

#### Ant assemblages

We used data from two previous studies in which ants were sampled systematically along two latitudinal gradients (Fig. 1). For the temperate-zone forests, Del Toro (2013) sampled ground-nesting ants at 67 sites spanning ten degrees of latitude and seven North American Level II ecoregions in the eastern United States. In the Neotropics, we focused on leaf-litter ant communities distributed throughout the Atlantic Forests of eastern Brazil. Leaf-litter ants are the most diverse ant fauna in the world in terms of phenotypic and functional diversity (Silva and Brandão 2010). The Atlantic Forest of eastern Brazil is the second largest tropical forest and spans the largest latitudinal gradient of all tropical forests in the Neotropics. All statistics analyses were carried out in the R v. 3.2.1 statistical environment (R Core Team 2015). The data and R code are archived online in the Harvard Forest Archives http://harvard-forest.fas.harvard.edu/data-archive.

**Figure 1.**
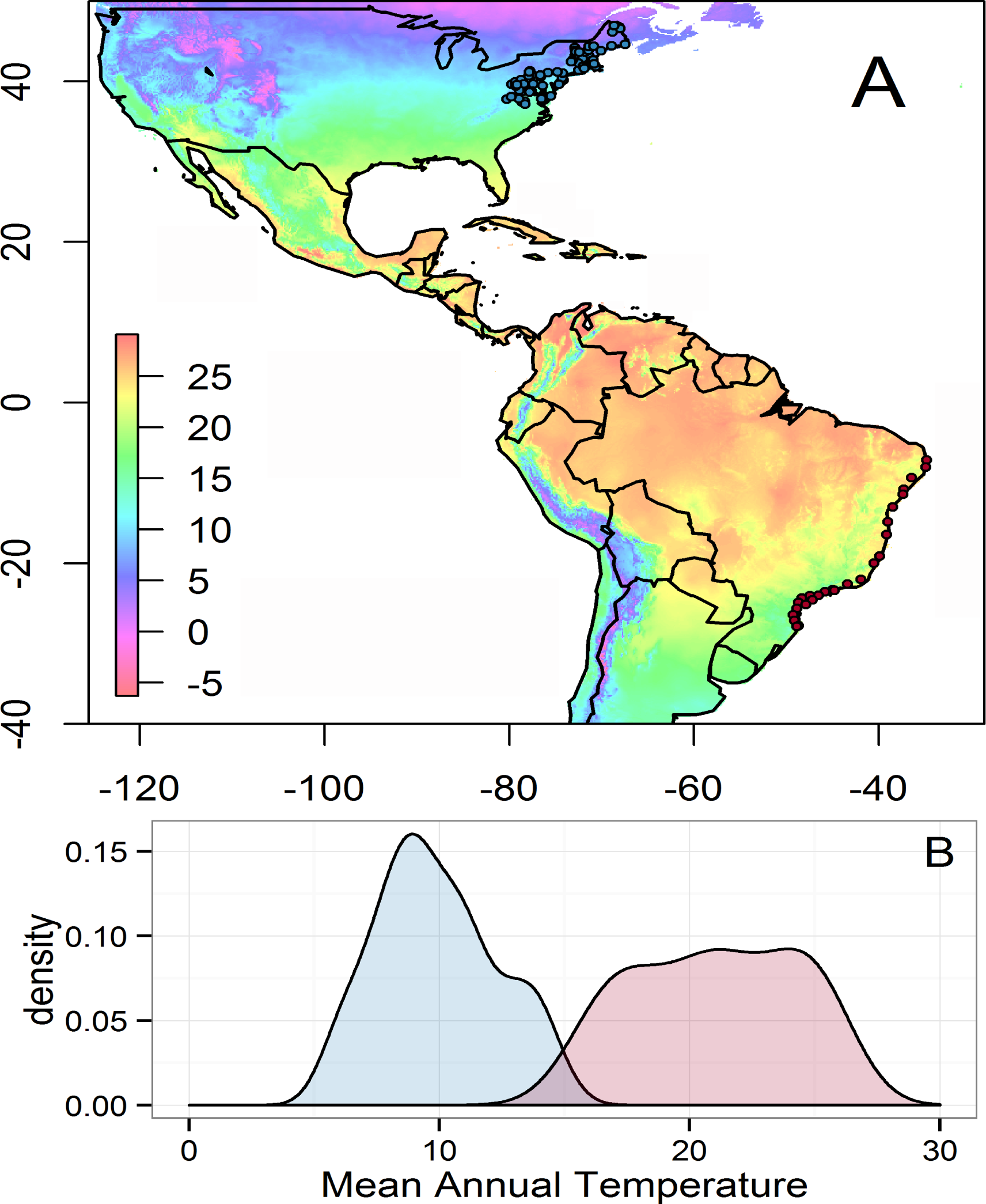
Study sites along two latitudinal gradients in temperate forests and in the South American Atlantic Forest (A), and frequency distributions of mean annual temperature in the regions (B).

### Morphological data

Our study draws on morphological data for 599 species of ants, 508 species from the Brazilian Atlantic Forest and 91 from temperate eastern North American forests. We examined fourteen characters that represent various aspects of ant ecology (Table 1; see Silva and Brandão 2010 for morphometric definition of each trait). These variables characterize morphological structures related to diet, habitat, and foraging strategies and are good proxy measures for ant species ecology (Silva and Brandão 2010, Arnan et al. 2014, Yates et al. 2014, Gibb et al. 2015b).

**Table 1.**
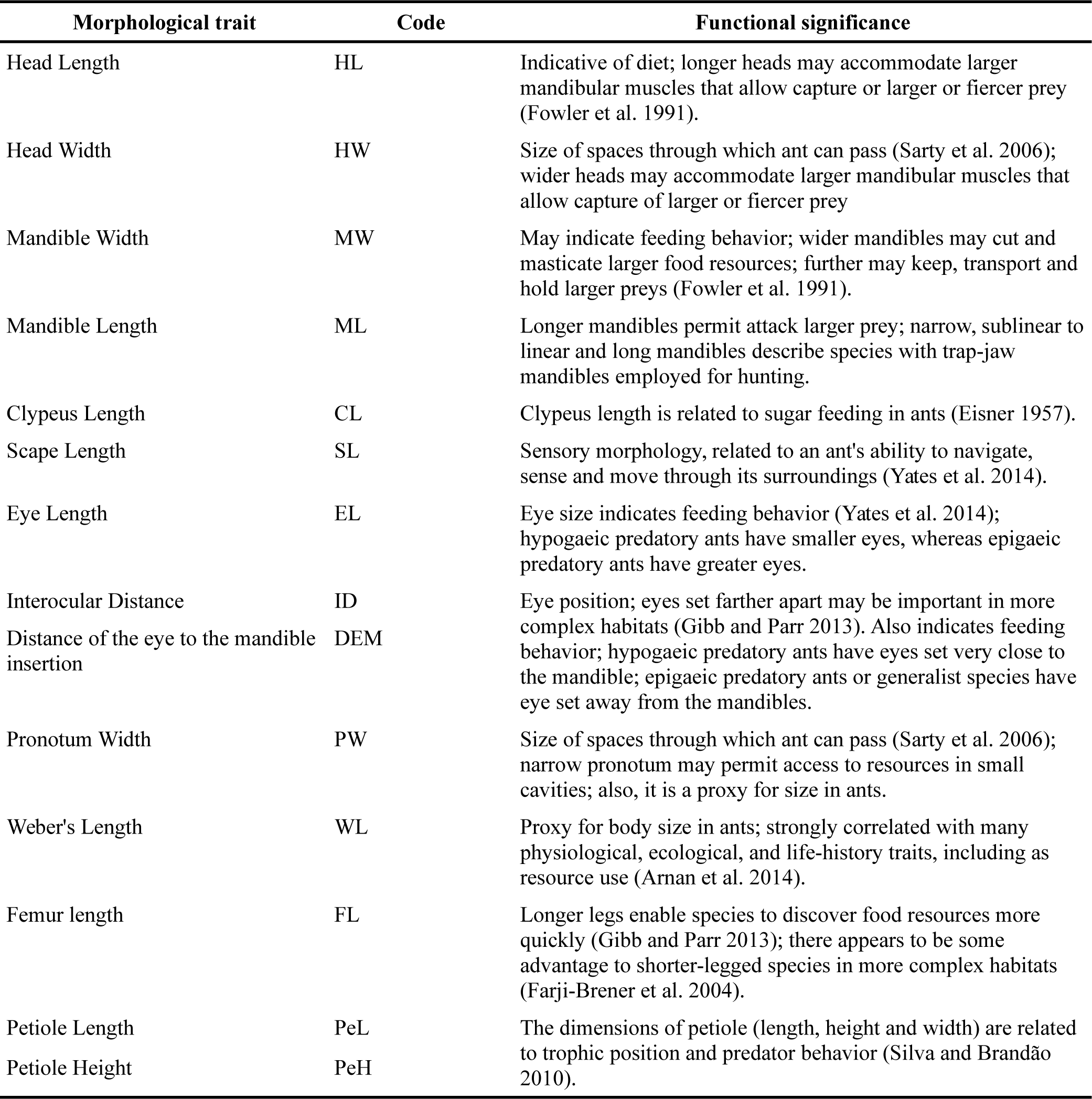
The fourteen ant traits used in this study, the code given to each trait and a description of the functional significance of traits.

Measurements of pinned specimens (± 0.01 mm) were made using an ocular micrometer attached to a Leica MZ75 stereomicroscope. We used the highest magnification that allowed the trait measured to be fitted within the range of the ocular micrometer. We measured fourteen morphological traits of between 1 and 25 (6 ± 4.7) individuals and of 1 to 11 (3 ± 1.4) individuals from the temperate and tropical datasets, respectively. A minimum of six individuals of each species were measured whenever possible. We used Weber's length (Table 1) as the descriptor of body size and used the other traits (head, mesosoma, metasoma, and body appendages) to describe the shape of each ant. For polymorphic or dimorphic species only measurements of minor-caste workers were included in the analyses.

We used log-transformed mean trait values for all analyses, as is routinely done for trait-based studies of continuous morphological characters (Trisos et al. 2014). Morphological traits were strongly and positively correlated (r = 0.65 – 0.96), largely because of their association with overall body size as well as phylogenetic correlations. To prevent these correlations from biasing analyses toward detecting only processes associated with body size, we used ordination techniques (based on principal component analysis) to derive independent trait axes.

### Statistical analyses

#### Principal component analysis

First we compared ant morphological diversity between the two biogeographic faunas (temperate in the Nearctic region and tropical in the Neotropical region), based on a principal component analysis calculated from the correlation matrix of the 14 log-transformed variables measured for all 599 ant species.

#### Morphological diversity metrics based on multiple traits

We used two standard metrics to quantify morphological diversity: functional diversity (FD: Petchey and Gaston 2002) and convex hull volume (CHV: Cornwell et al. 2006). Morphological diversity is a measure of how dispersed a set of species is in the trait space (Petchey and Gaston, 2002), while CHV is the smallest convex set in trait space enclosing all species trait values within a community (Cornwell et al. 2006). We selected FD and CHV because they are widely used with presence/absence data and have been shown to have a relatively high power to detect trait clustering (CHV) and overdispersion (FD) in community assembly simulations (Mouchet et al. 2010, Aiba et al. 2013). We calculated FD and CHV using three derived trait principal axes that were centered and relative to the species pool (i.e., the full data set of trait values for each region; see also Villéger et al. 2008). Because they test concurrently for habitat filtering and niche differentiation, both FD and CHV are considered to be multiple-niche-axis metrics (Trisos et al. 2014).

We also estimated mean morphological diversity (MPD) and mean nearest-neighbor distance (MNND), both of which have been used commonly as measures of community relatedness and have been the subject of previous power analyses (Kraft et al. 2007, Trisos et al. 2014) (Table 2). MPD is the mean of pairwise distances between co-occurring species and is most sensitive to tree-wide patterns of morphological clustering and evenness (Trisos et al. 2014). Mean pairwise-distance measures the volume of all species in a community within the morphological space (Webb 2000). MNND is the mean of the morphological distances separating each species from its closest cooccurring relative and is most sensitive to patterns of morphological clustering or evenness at the tips from a cluster dendrogram of morphological similarity (Kraft et al. 2007); MNND measures how clumped species are in morphological space (Webb 2000).

**Table 2.**
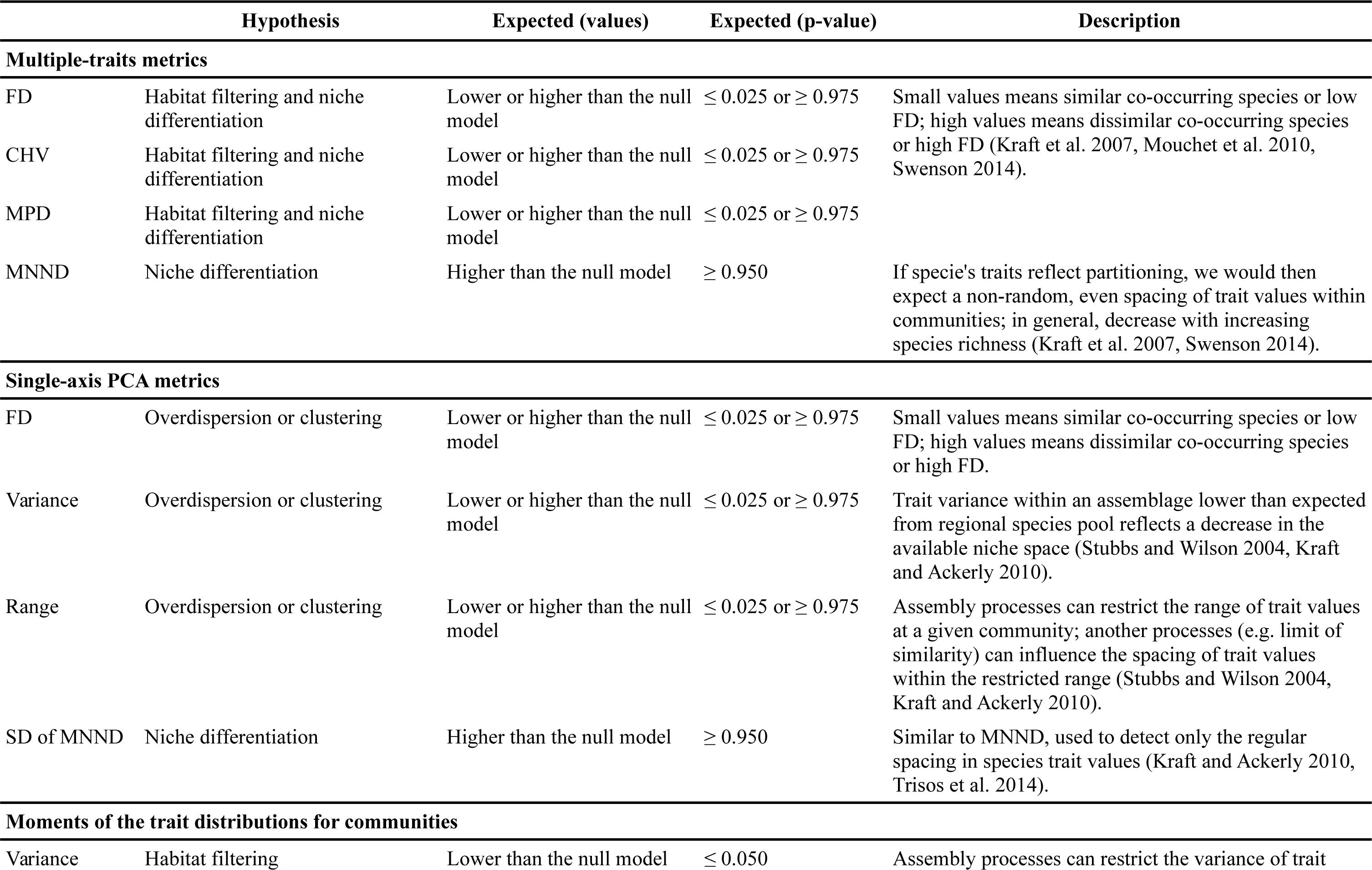
List of selected morphological metrics, hypothesis tested, predictions, and description of the biological significance.

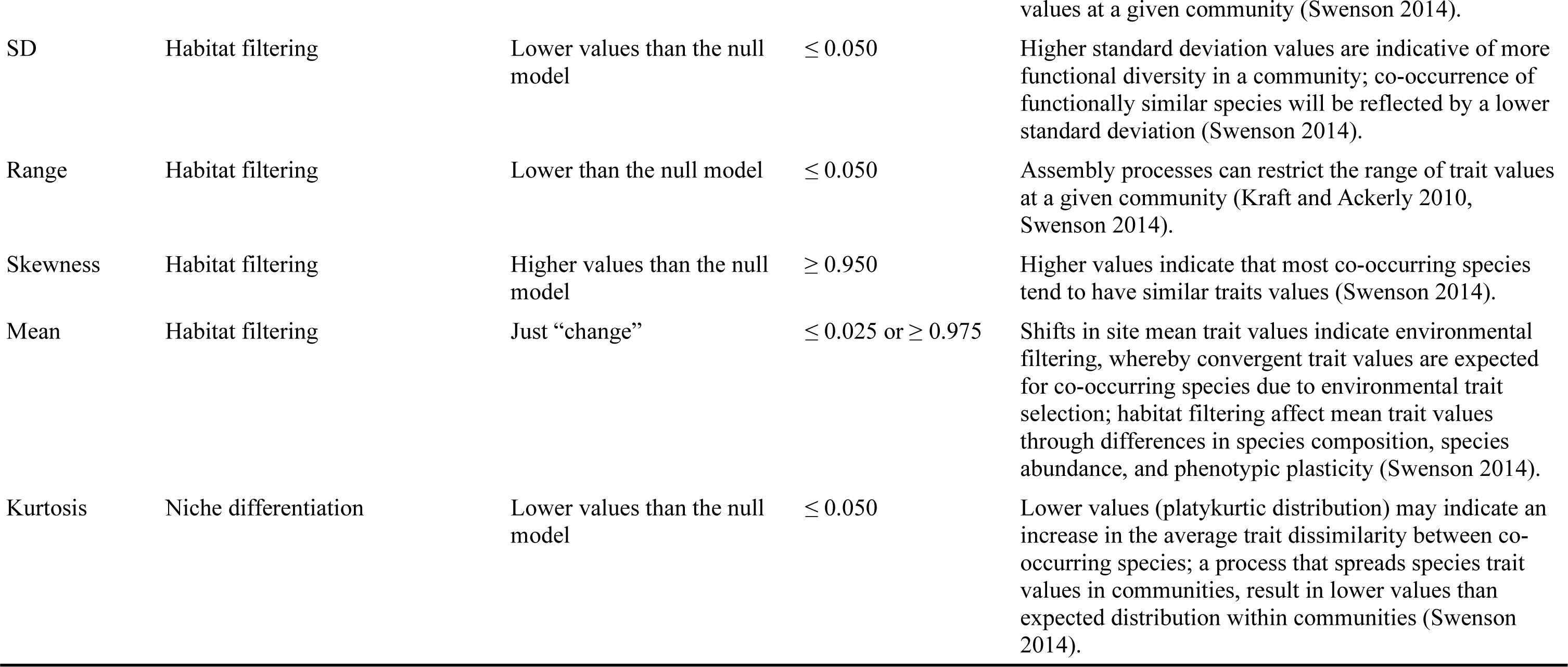

Further, we analyzed each derived trait axis individually using metrics to test for both assembly patterns (clustering and overdispersion) and metrics used to test for only one assembly pattern (Table 2). We focused on three multi-pattern metrics (Trisos et al. 2014): (1) FD applied to a single trait axis; (2) variance in species values within an assemblage along a single trait axis; and (3) range in species trait values within an assemblage. Variance and range have also been used as measures of trait clustering (Stubbs and Wilson 2004, Kraft et al. 2008, Kraft and Ackerly 2010). We also used one single pattern metric, SD of MNND (the standard deviation of the distances between neighboring species along a single trait axis). This latter index quantifies evenness of species dispersion or packing in morphological space, and is used only to detect the regular spacing in species trait values predicted by competitive exclusion (Kraft and Ackerly 2010). We calculated these metrics using two derived trait principal axes that were centered and relative to the species pool (i.e., the data set of trait values for each regional species pools separately).

We tested differences in morphological diversity between tropical and temperate ant assemblages using linear mixed effect models (LMEs) with a random intercept and random slope terms. We compared models with the same random effect structure but different fixed structure (i.e., morphological diversity ~ observed species richness or morphological diversity ~ observed species richness + region); the random structure was observed species given region. The LMEs allow us to compare morphological diversity between regions accounting for species richness differences in the ant assemblages. We used the *lmer* function in nlme version 3.1-128 (Pinheiro et al. 2016) to build the models.

#### Morphological diversity metrics based on single traits

We also selected eight “priority” ant traits to calculate community metrics of spacing of trait values (range, variance, SD, mean, kurtosis and skewness), to explore deterministic process in the communities. For example, if habitat filtering occurred at the sampling scale, the range of observed trait values would be smaller than that expected under a null model. Whereas the range is useful to detect effects of a habitat filter, it does have the statistical downside of being susceptible to extreme observations that could be due to mass effects (Shmida and Wilson 1985). For this reason, it is important to present results for both range and variance of the traits (Cornwell and Ackerly 2009). Note that habitat filtering may reduce variance but it may also be affected by niche differentiation. Habitat filtering restricts the range of trait values and the limits of similarity affect spacing (Cornwell and Ackerly 2009); both of these processes are likely to affect the variance of trait values among species within a community. Habitat filtering may shift the mean of the trait distribution relative to a null expectation (but also can occur without this effect; Kraft et al. 2008). Likewise, if niche differentiation is occurring, the kurtosis of the distribution of trait values, will be smaller than expected from a null model (Stubbs and Wilson 2004, Cornwell and Ackerly 2009).

We calculated the variance, SD, range, mean, skewness, and kurtosis for the eight selected ant traits (HL, SL, EL, MW, DEM, FL, ID, PeH) based on presence-absence of species at a site because we were interested in quantity changes that arise from a replacement of species with different traits, which reflects among-species variation within communities (Cadotte et al. 2010).

#### Null models

We used null models to generate expected values of community structure given the observed number of species present at each site. We used null models to help identify patterns in the assemblages rather than to identify specific ecological processes, including those that alter dispersion of a trait (or traits) within a species assemblage as a filter (biotic or abiotic). In the null models, both number of species and occurrence of species in the assemblages were maintained. Null model simulations were conducted in R using functions from Swenson (2014) modified for our datasets. A measure of structure for a particular assemblage was considered significant if it fell outside the extremes of the distribution of random structures generated from the null model (Table 2).

We defined different species pools to assemble the null communities, constraining which species could disperse to a given location in the tropical and temperate regions. For the tropical data set, we used the Atlantic Forest regions defined by Silva and Brandão (2014) and for the temperate data set we used the five different EPA Level II ecoregions (Del Toro 2013). The following regions were used in the analysis: temperate data set: (1) Appalachian Forests (18 sites and 63 species), (2) Atlantic Highlands (22 sites and 65 species), (3) Mixed Wood Plains (13 sites and 53 species), (4) Southeastern Coastal Plains (7 sites and 54 species), (7) Southeastern US Plains (7 sites and 44 species); tropical data set: (1) high southeastern-south areas (8 sites, 262 species), (2) low southeastern-south areas (6 sites, 229 species), (3) intermediate latitude areas (4 sites, 239 species), and (4) north Atlantic Forest areas northeastern (8 sites, 233 species).

#### Environmental gradients and morphological traits

Geographic distributions of ants are affected by a number of environmental characteristics; temperature and precipitation are usually the strongest direct and indirect correlates (Dunn et al. 2009, Jenkins et al. 2011, Gibb et al. 2015a) as they possibly affect the amount and distribution of available resources. We therefore tested the hypothesis that aspects of temperature variability and precipitation were important predictors of ant morphological structure. We obtained for each site estimates of precipitation and minimum temperature values based on data from WorldClim climate data layers (Hijmans et al. 2005). Annual precipitation (bioclimatic variable 12) is defined as the sum of all the monthly precipitation estimates; minimum temperature of coldest month (bioclimatic variable 6) is defined as the lowest temperature of any weekly minimum temperature.

First, we used Pearson's correlation coefficients to examine the relationships between means of trait values and the two climatic variables. Then, we tested the relationship between the eight selected traits and the two climatic variables. Relationships between trait means and variances in assemblages and climatic variables can be complex, and may exhibit interactions between drivers and non-linear responses (Símová et al. 2015). We examined these complex relationships using multiple regression in which the eight trait values entered as response variables, while all the environmental variables (linear and quadratic terms) and their interaction terms entered the model as driver variables. Because there is a low overlap in climate between tropical and temperate zones, observed relationships could arise solely because of mean trait differences between zones, and without any actual relationship between traits and climate within each zone. Thus, we included region as a term in the model. We used stepwise selection by AIC in the *MASS* package (Venables and Ripley 2002) to determine the best of the multiple regression models. We used a Type III ANOVA to test for the presence of a main effect after model selection using the *anova*function in the *car* package (Fox and Weisberg 2011), as this approach does not depend on the order of explanatory variables in the model.

## Results

### The morphospace of temperate and tropical assemblages

The first principal component based on the total dataset represented a general size axis accounting for 79% of the total variance, followed by component 2 (9%, eye size and petiole height) and component 3 (3%, distance of eyes to the mandibles and mandible width) (Fig. 2, Supplementary material Appendix 1, Table A1).

**Figure 2.**
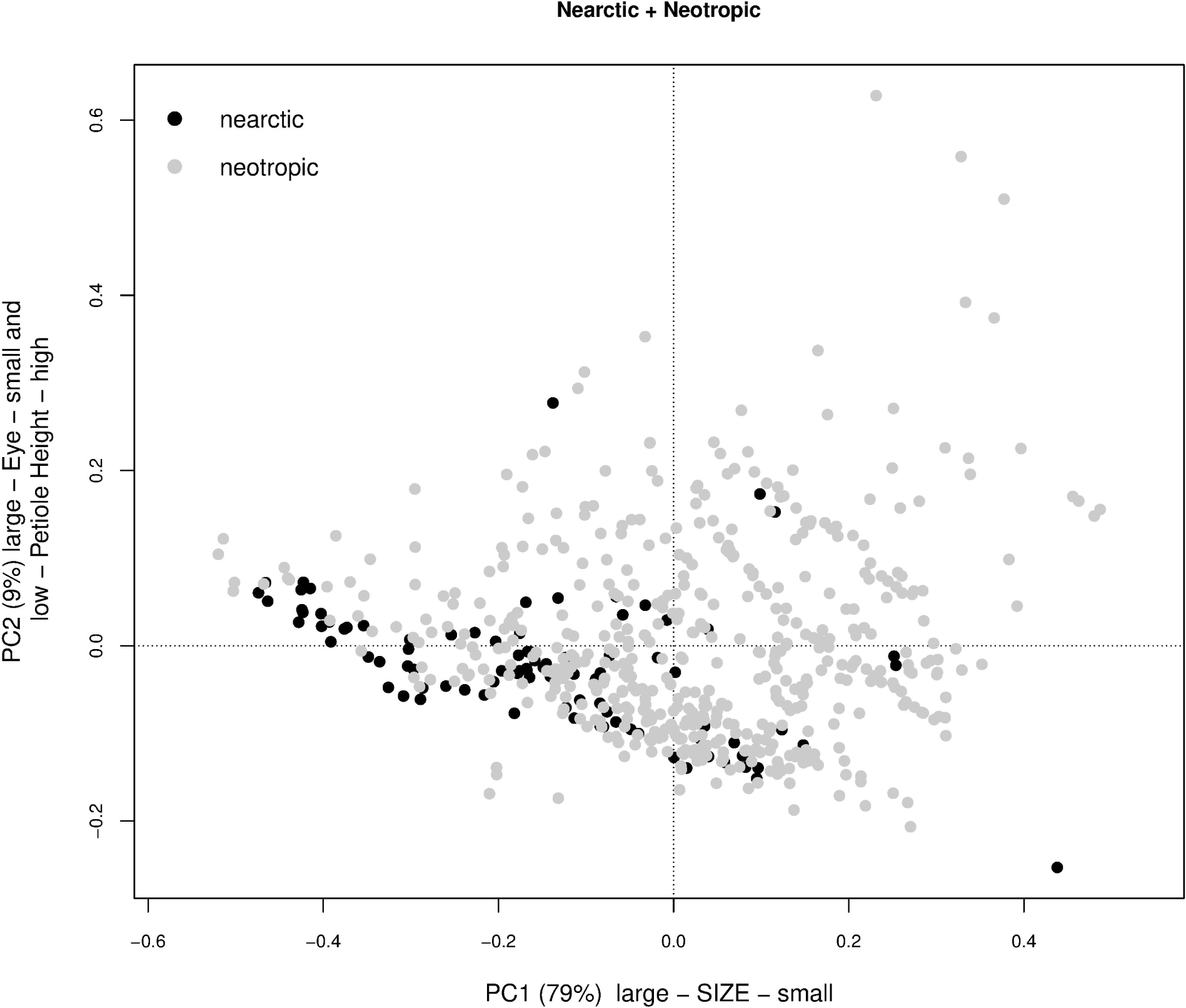
The morphological distribution on one size and one shape principal component axis for tropical (light gray dots) and temperate-zone forest (black dots) ants morphospace (N= 599).

### Trait analyses

All results of trait analyses were based on constrained species pools to quantify trait diversity in the ant assemblages. However, unconstrained analyses produced similar results. Tropical assemblages had higher morphological diversity and morphological volume than temperate assemblages (Fig. 3a-b); however, conditional on species richness, MPD and MNND values were not higher in tropical forests than temperate forests (Fig. 3c-d; Supplementary material Appendix 1, Table A2; Appendix 2, Figs. A1-A2). Temperate assemblages had larger variance in size (PC1) whereas tropical assemblages had larger range values. Functional diversity metric (FD) from PC1 axis had higher values in tropical assemblages. SD of MNND suggested higher species packing in tropical assemblages as calculated from PC1 axis (Fig. 4a-d). However, the differences in trait diversity values in tropical assemblages as calculated from PC1 axis were not higher than temperate forests given their species richness (Supplementary material Appendix 1, Table A2).

**Figure 3.**
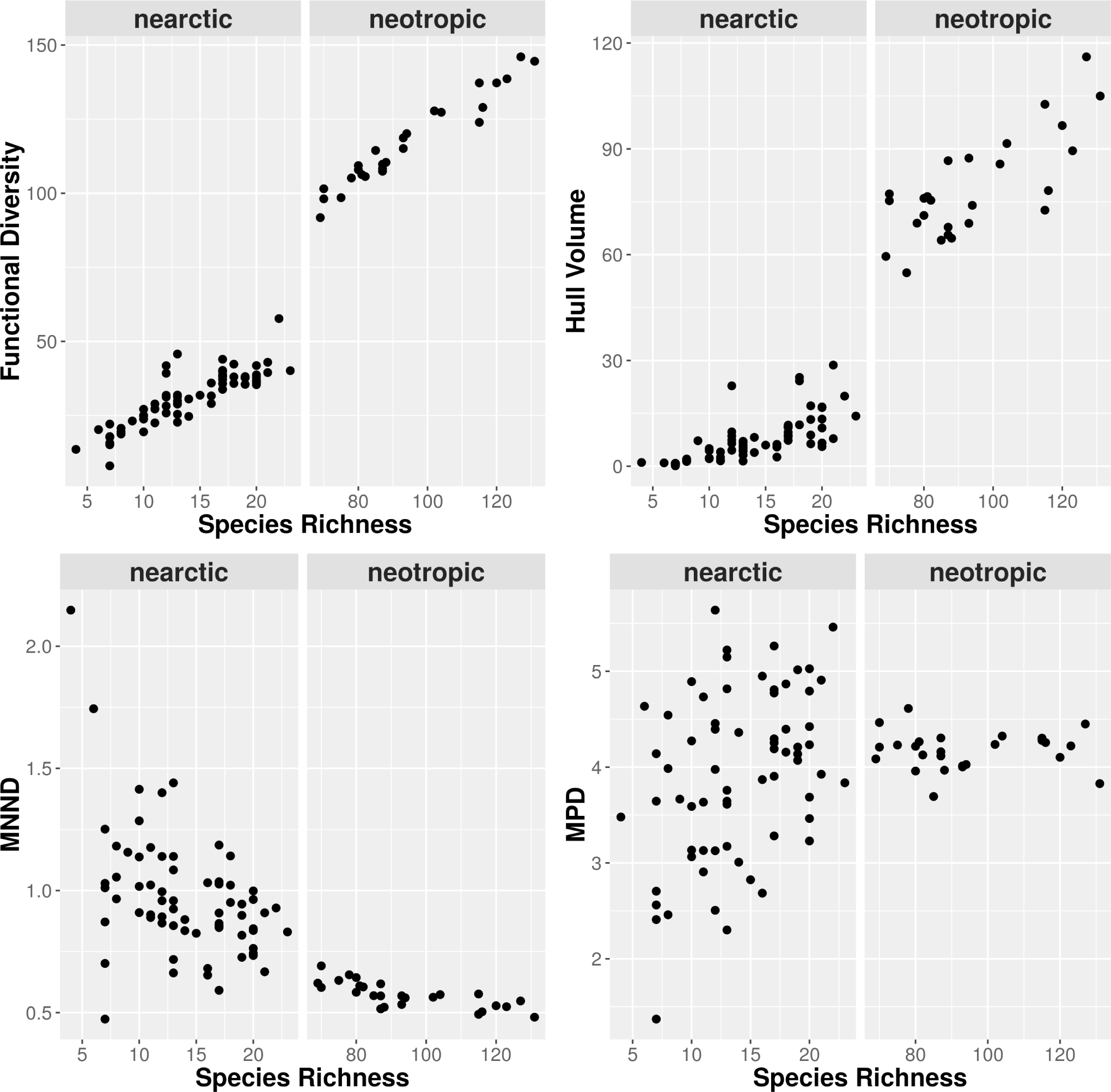
Four measures of distance between species (multiple-axis metrics) in the morphospace plotted as functions of the number of species in the ant assemblages in the temperate and tropical regions: (a) Morphological Diversity (FD), (b) convex hull volume (CHV), (c) mean morphological distance (MPD), and (d) mean nearest morphological distance (MNND). Each metrics was calculated using species scores on the first three principal component axes, derived from constrained species pools for each region separately; each species is a composite of fourteen traits.

**Figure 4.**
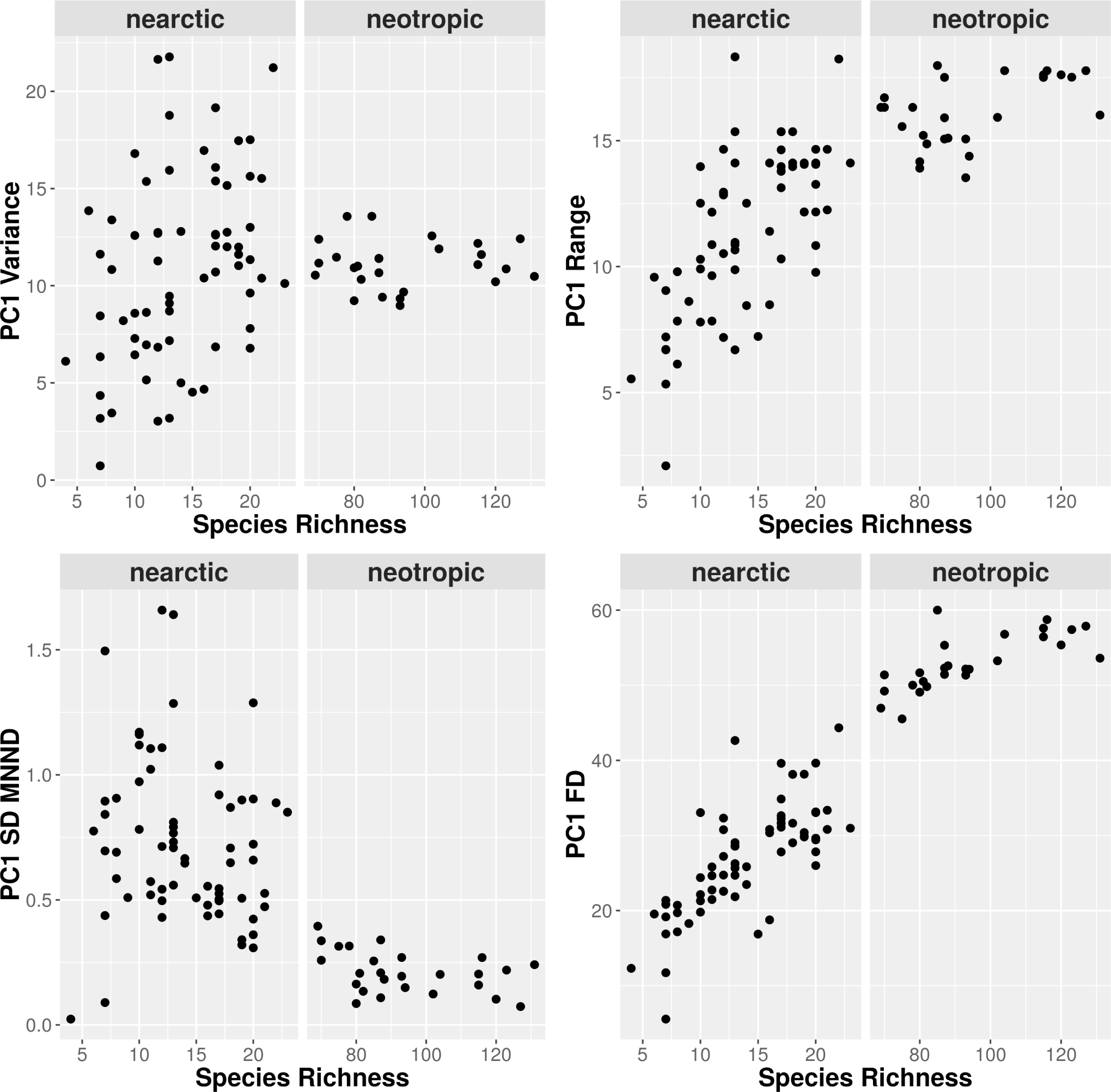
Four measures of distance between species (single-axis metrics) in the morphospace plotted as functions of the number of species in the ant assemblages in the temperate and tropical regions. (a) Variance, (b) Range, (c) standard deviation of MNND (SD of MNND), and (d) Functional Diversity (FD). Each metrics was calculated using PC 1 axis, derived from constrained species pools for each region separately; each species is a composite of fourteen traits.

### Single trait analysis

The moments of trait distributions indicated clear trait differences between temperate and tropical assemblages. The community mean of traits was strongly different between regions; temperate assemblages had larger values of traits and tropical assemblages higher density of small values (Fig. 5). Further, tropical assemblages had higher asymmetry in mean trait values. On the other hand, the distribution of mean traits calculated from species pools had large overlap between regions (Supplementary material Appendix 2, Fig. A3).

**Figure 5.**
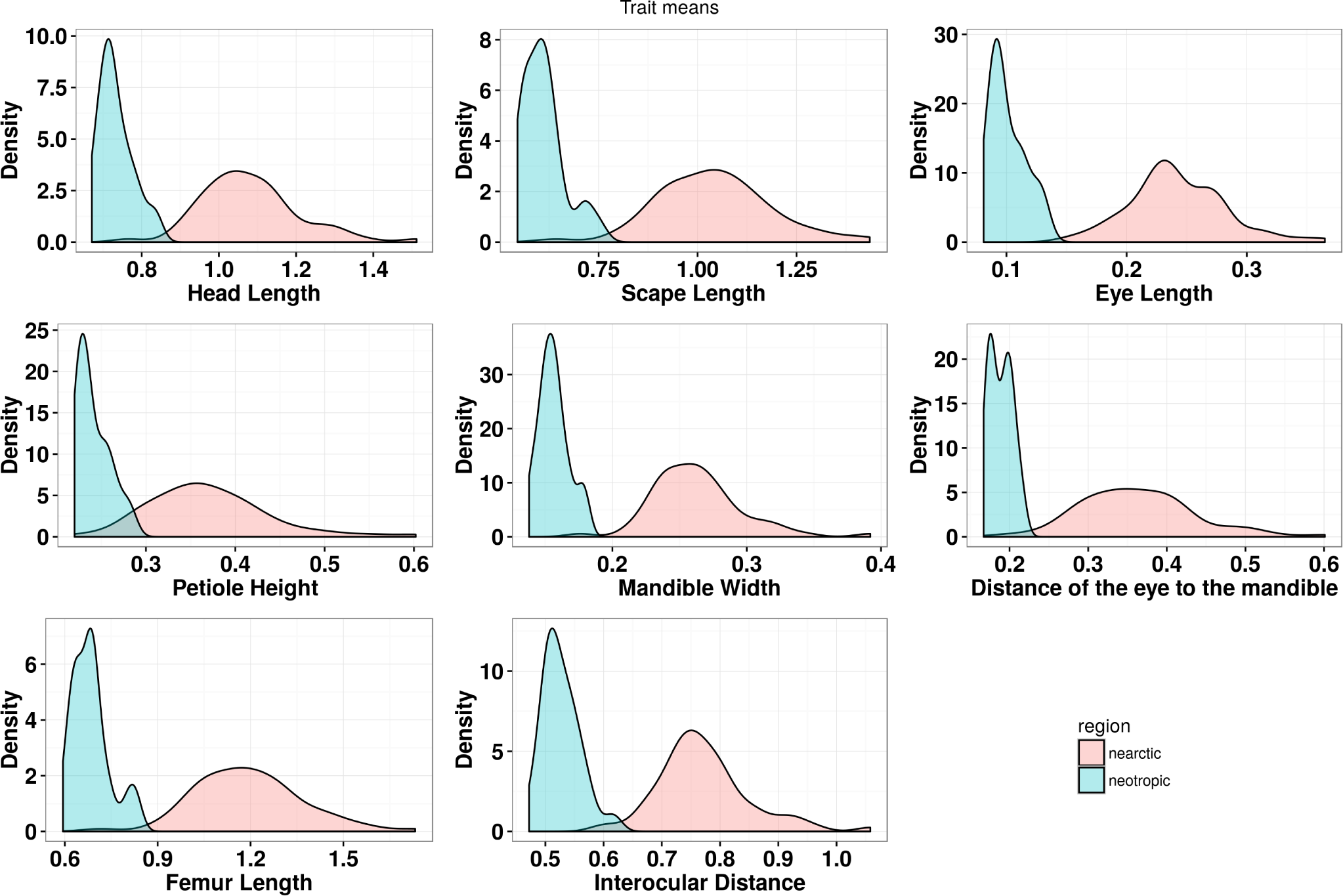
Frequency distributions of trait means at assemblage scale in the temperate and tropical regions.

Neotropical assemblages had higher right-skewed trait distribution suggesting that most cooccurring species tended to have similar trait values (as also suggested by MNND and SD of MNND) (Supplementary material Appendix 2, Fig. A4). Distance of the eye to the mandibles had larger overlap in skewness distribution at the assemblage level. However, the distribution of community variance was relatively similar between regions, except for the variance of the distance of the eye to the mandibles in the temperate region (Supplementary material Appendix 2, Fig. A5).

Kurtosis had strong differences between regions. In general, temperate assemblages had smaller values, suggesting more platykurtic distribution and indicating an increase in the average trait dissimilarity between co-occurring species (Supplementary material Appendix 2, Fig. A6). Finally, we did not find clear differences in trait ranges; scape length, eye length and the distance of the eye to the mandibles had large overlaps in range of values (Supplementary material Appendix 2, Fig. A7).

### Null model analysis

Overall, we found weak evidence of habitat filtering or niche differentiation at the assemblage scale. Non-random values in the morphological metrics were rarely observed. The proportion of temperate assemblages in which we detected significant values in the metrics ranged from less than 1 to 16% and was rarely observed in the Neotropical region. Habitat filtering was detected more commonly in the temperate assemblages, especially for the moments of trait distributions for eight selected ant traits (Supplementary material Appendix 1, Table A3). This pattern also was detected for single trait analysis (PC1 or PC2), although significant results decreased when PC2 was considered (probably because PC2 had a very low contribution to the variance in the morphospace). Significant lower morphological diversity than expected was detected mainly in temperate assemblages (Supplementary material Appendix 1, Table A3, for multiple trait metrics). Null models testing using constrained species pools yielded similar deviation from the null expectation, consistent with the results of unconstrained analysis (Supplementary material Appendix 1, Table A4).

### The relationship between assemblage trait variation and climate

Mean trait values of all 599 measured ant species were characterized by high correlations, which influenced the relationship between trait means and climate variables. Trait means of HL, SL, EL, PeH, DEM, and FL all were best explained by the region term in the model, without climate relationships; tropical assemblages were characterized by lower mean trait values than temperate assemblages. By the other hand, MW and ID means were best explained by the quadratic term of temperature and region; the relationship was concave and trait means values decreased with increasing temperature (Supplementary material Appendix 1, Table A5). The relationship between trait variances and climate were more variable, suggesting that variation in variances in trait values was driven by region and multiple environmental factors. Trait variances of EL, PeH, MW, and the DEM were all best explained by region, without climate relationships; temperate assemblages had larger variance in traits than tropical assemblages. Variance in HL and FL were best explained by the quadratic term of temperature and region; the relationship was concave and variance values decreased with increasing temperature. The quadratic term of precipitation, region and the interaction term between temperature and precipitation were the best predictors of variance in SL. Finally, ID had no relationship with region or climate (Supplementary material Appendix 1, Table A5).

## Discussion

We found strong differences in morphological space occupied by ant communities in temperate and tropical regions. As expected, ant communities in tropical areas showed higher morphological diversity; however, temperate ant communities showed higher variance in size. We found evidence that tropical assemblages harbor higher levels of morphological diversity that the temperate-zone ants. In general, more diverse niches and fewer gaps within the total niche space are expected in diverse tropical communities than in those from similar habitats containing fewer species at higher latitudes (Winemiller et al. 2015).

Although there was an overall higher diversity of traits in the tropics, we unexpectedly did not find more tightly packed species in the tropics within the trait space. Such species packing could result from a lack of biotic interactions (or Connell's “ghost of competition past”), low population sizes reducing the encounter rate of species, or finer partitioning of local scale environmental gradients (Swenson and Weiser 2014). It is interesting that different results (higher functional diversity and species packing) were found in comparisons of plant functional diversity between North and South America (Swenson et al. 2012). Recent studies on phylogenetic structure of arboreal (Blaimer et al. 2015) and leaf-litter ant communities (Donoso 2014) also have suggested that habitat filtering structures the communities in humid forests, selecting for related species with similar ecological traits and leading to a clustered pattern. By contrast, in temperate forests of North America and Europa, Machac et al. (2011) found evidence either for overdispersion or random structure at lower elevation and clustering at high elevation sites.

Because temperate-zone ant communities showed similar species packing (quantified as the mean nearest neighbor distance in multivariate trait space), we hypothesize that the higher morphospace volume and functional diversity in tropical communities occurs via an increase in the morphological volume. This also suggests that as species are added to assemblages they tend to expand the volume to a greater degree than they pack the functional volume.

### Null models and morphological structure

There was little evidence within either temperate or Neotropical ant assemblages of habitat filtering or niche differentiation patterns based on our null models, although as expected, habitat filtering was detected more commonly in the temperate-zone ant assemblages. In abiotically harsh environments assemblages should contain a non-random subset of species that are more functionally similar than expected (Swenson et al. 2012). The absence of both clustering and overdispersion in traits has been reported fairly widely in plants and for some animal communities at local scales, and is typically interpreted as evidence for a neutral model of community assembly (Trisos et al. 2014). The use of multiple niche axis metrics, as mean pairwise morphological distances (a measure of overall dispersion of morphology for each local assemblage), detected that 16% of temperate assemblages had lower than expected functional diversity given their observed species richness. In fact, niche filtering rather than niche partitioning appears to structure ant assemblages in temperate forests in the south-eastern United States (Fowler et al. 2014). It is relatively well known that processes that shape ant communities depend on spatial scale and likely vary among ecosystems (Nipperess and Beattie 2004, Gotelli and Ellison 2002, Sanders et al. 2007, Fowler et al. 2014).

We found weak patterns of niche differentiation in the tropical dataset. Community ecology studies generally predict that niche differentiation will be a dominant influence on community assembly in the tropics (Algar et al. 2011, Winemiller et al. 2015). Our null model analysis may be potentially biased because the analyzed scale may be inadequate to detect niche differentiation evidences. However, similar analyses for the tropical data set by Silva and Brandão (2014) using a smaller number of traits and constrained pool null models also did not find strong evidences of under- or overdispersion in the leaf-litter ant fauna, but suggested shifts in the morphospace structure along the latitudinal gradient. In this case, co-occurrence analysis at sample point scale (1 m2) or constrained by guilds can be particularly informative (Silva and Brandão 2014) because patterns of morphological structure should be more probable within, rather than across feeding guilds.

The prediction that under more harsh climatic conditions the North America assemblages should be constrained more often by habitat filtering also was supported only weakly by examination of a single niche axis. Our result appear to contradict recent studies on habitat filtering that found strong significant evidence for convergence in ant communities traits in North America (e.g., Fowler et al. 2014). In conclusion, we did not find strong pattern of non-random convergence or divergence in traits when comparing to the species pool of temperate or tropical regions. Rather, the link between ecological assembly processes and trait patterns might be complex and weak (Gerhold et al. 2015).

### Single trait analysis

The use of selected single traits to compare trait distribution between assemblages was useful in describing differences between regions. We suggest that focused analysis of single traits may be an important approach to compare and describe the distribution of ant functional diversity. Trait-based metrics combine multiple traits into a single analysis (e.g., functional diversity) and because different traits are often associated with different niche axes, these metrics may have the advantage of providing an integrated overview of community structure (Trisos et al. 2014). However, if different assembly processes act on separate traits or exert combined effects, a potential drawback is that multiple-traits may combine signals of contrasting assembly processes (Swenson and Enquist 2009, Trisos et al. 2014).

The moments of trait distributions showed marked differences between temperate and tropical assemblages. For example, the distribution in size of ant attributes was strongly right skewed, indicating higher density of smallest ants in tropical sites. The bulk of leaf-litter ant species is composed by very small species, spanning diverse trophic groups (generalist and specialist predators, omnivorous species, from small to medium size; Silva and Brandão 2010, Donoso, 2014).

Single trait analyses revealed not only differences between ant morphology in tropical and temperate assemblages, but also different within-region responses of traits in variance, kurtosis, skewness and range. Although there was a high correlation between traits, the analysis of the moments of trait distributions suggested that all selected traits can be important to detect differences in morphological structure. In particular, both variance and kurtosis in the distance of the eye to the mandible insertion showed it to be the strongest difference comparing assemblages and we suggest that it may be an important attribute to trait-based analysis of ant communities. Eye morphology seems to be particularly important in trait-based studies; eye relative size in ants is related to period of activity (Yilmaz et al. 2014), degree of predatory behavior (Fowler et al. 1991), determines the ability to see laterally, and influences the success of species travelling through complex habitats (e.g., leaf-litter habitat) (Gibb and Parr 2013).

The trait distributions in skewness or kurtosis also suggest that in Neotropical ant assemblages most co-occurring species tend to have similar traits, whereas in temperate assemblages there tends to be on average higher trait dissimilarity between co-occurring species. In fact, the results of multiple trait analysis suggest that temperate communities have higher trait dissimilarity than tropical communities. Of course, tropical species exhibit an extraordinary variety of adaptations molded by their interactions with resources, competitors, and antagonists (Ricklefs 2012). In sum, our results suggest that the larger tropical species pools occupy a morphological space distinct from that of temperate species but not a more tightly packed morphospace.

In general, we can confidently describe that leaf-litter ant fauna in the Neotropics as composed by more specialist taxa (Delabie et al. 2000) and that the epigaeic temperate-zone ant fauna includes predominantly opportunistic and omnivorous feeders species (Ellison et al. 2013). The range distribution of a petiole measure, as such petiole height, showed strong difference between tropical and temperate ant assemblages. Ant petiole measures may be used to describe morphological groups of species, especially for predator species (Silva and Brandão 2014). We suggest that petiole measures also should be considered to understand how ant traits respond to environment, especially in the tropics where predators are often an important component of communities.

Taken together, the results of single trait analysis suggest that choosing traits that can be used to test hypotheses can be important for detecting differences in morphological structure of ant communities, especially when considering effects of disturbance and climatic change on assemblage structure. Our results also support a recent trend in community ecology that proposes not only the integration of data sets but also their subdivision into niche axes before analysis (Spasojevic and Suding 2012, Trisos et al. 2014, Winemiller et al. 2015).

### Ant traits and habitat characteristics

Ant traits have been shown to respond to habitat characteristics such as structural complexity (Farji-Brener et al. 2004, Gibb and Parr 2010, Yates et al. 2014). For example, leg length decreases with habitat complexity (Farji-Brener et al. 2004, Sarty et al. 2006, Gibb and Parr 2010). Gibb and Parr (2013) extended analysis testing morphological trait responses to habitat complexity in sites on three continents, finding predictive responses of sensory morphology of ant assemblages to habitat complexity. Eye position has been suggested as a key response variable to predatory behavior (Silva and Brandão 2014) and habitat complexity (Gibb and Parr 2013). Clear differences between macrohabitats in in ant's scape length, eye size, leg length, and body size have been described (Yates et al. 2014, Schofield et al. 2016); further, ant morphology may respond to competition (Nipperess and Beattie 2004).

Temperature and precipitation are considered drivers of ant species richness patterns at global scale (Dunn et al. 2009, Jenkins et al. 2011, Gibb et al. 2015a) and recent studies also suggest a strong relationship between temperature and ant functional responses (Diamond et al. 2012, Stuble et al. 2013). Ant functional diversity decreased with reduced temperature in a high elevation gradient (Raymond et al. 2013) and climate seasonality played an important role in shaping the occurrence of functional species traits in Mediterranean ant communities (Arnan et al. 2014). Our analysis suggests that trait mean values all were correlated with region; size also decreased with increasing temperature for some traits. Our environmental model largely reflects the smaller ants in tropical assemblages. On the other hand, the environmental model for trait variances largely reflects the higher variance in ant traits in temperate assemblages. However, the response of trait variances was more complex and the explained variance by models was lowest compared to the trait mean model. For example, head length, and femur length correlated negatively with temperature. Interactions between temperature and precipitation were detected for trait related to food search (as scape length), suggesting that relationships between climate and traits varies for ants among regions.

### Concluding remarks

The human-induced extinction of species in the Anthropocene (Dirzo et al. 2014) with ongoing biotic impoverishment may alter key ecosystem processes with important consequences for ecosystem services needed by the humanity (Mouillot et al. 2014). Our results indicate that ant traits respond to environmental gradients and single trait analysis may be an important approach to analyze the loss of particular functions delivered by ants. This approach may guide future studies on ant responses to habitat modification and climate change. Although many aspects of the morphological structure in ant communities are as yet unexplored (including the relationship between morphology and the variety of resources), our results underline the importance of morphological analyses in arthropod communities and point out to the need for more through characterization of morphological space for modeling the impacts of habitat disturbance and climate change on morphological diversity.

## Acknowledgments

Funding was provided by grants from the Fundação de Amparo à Pesquisa do Estado de São Paulo (2010/51194-1 and 2010/20570-8), and by Conselho Nacional de Desenvolvimento Científico e Tecnológico (CNPq) through a PCI short-term grant (170040/2015-1) to Rogério R. Silva.

## References

Aiba, M. et al. 2013. Robustness of trait distribution metrics for community assembly studies under the uncertainties of assembly processes. – Ecology 94: 2873–2885.

Algar, A. C. et al. 2011. Quantifying the importance of regional and local filters for community trait structure in tropical and temperate zones. – Ecology 92: 903–914.

Arnan, X. et al. 2014. Ant functional responses along environmental gradients. – J. Anim. Ecol. 83: 1398–1408.

Blaimer, B. B. et al. 2015. Functional and phylogenetic approaches reveal the evolution of diversity in a hyper diverse biota. – Ecography 38: 901–912.

Cadotte, M. W., et al. 2010. Phylogenetic diversity metrics for ecological communities: integrating species richness, abundance and evolutionary history. – Ecol. Lett. 13: 96–105.

Cadotte, M. W. et al. 2013. The ecology of differences: assessing community assembly with trait and evolutionary distances. – Ecol. Lett. 16: 1234–1244.

Cadotte, M. W. et al. 2015. Predicting communities from functional traits. – Trends Ecol. Evol. 30: 510–511.

Cornwell, W. K. and Ackerly, D. D. 2009. Community assembly and shifts in plant trait distributions across an environmental gradient in coastal California. – Ecol. Monogr. 79: 109–126.

Cornwell, W. K. et al. 2006. A trait-based test for habitat filtering: convex hull volume. – Ecology 87: 1465–1471.

Delabie, J. H. C. et al.. 2000. Litter ant communities of the Brazilian Atlantic rain forest region. In: Agosti, D., Majer, J.D., Alonso, L.E. and Schultz, T. Sampling ground-dwelling ants: case studies from world’s rain forests. – Curtin University School of Environmental Biology Bulletin 18: 1–17.

Del Toro, I. 2013. Diversity of Eastern North American ant communities along environmental gradients. – PLoS One 8: e67973.

Del Toro I. et al. 2012. The little things that run the world revisited: a review of ant-mediated ecosystem services and disservices (Hymenoptera: Formicidae). – Myrmecol. News 17: 133–146.

Del Toro, I. et al. 2015. Ant-mediated ecosystem functions on a warmer planet: effects on soil movement, decomposition and nutrient cycling. – J. Anim. Ecol. 84: 1233–1241.

Diamond, S. E. et al. 2012. A physiological trait-based approach to predicting the responses of species to experimental climate warming. – Ecology 93: 2305–2312.

Dirzo R. et al. 2014. Defaunation in the Anthropocene. – Science 345: 401–406.

Donoso, D.A. 2014. Assembly mechanisms shaping tropical litter ant communities. – Ecography 37: 490–499.

Dunn, R. R. et al. 2009. Climatic drivers of hemispheric asymmetry in global patterns of ant species richness. – Ecol. Lett. 12: 324–333.

Eisner, T. 1957. A comparative morphological study of the proventriculus of ants (Hymenoptera: Formicidae). – Bull. Mus. Comp. Zool. 116: 441–490.

Ellison, A. M. et al. 2012. A field guide to the ants of New England. Yale University Press.

Farji-Brener, A. G. et al. 2004. Environmental rugosity, body size and access to food: a test of the size-grain hypothesis in tropical litter ants. – Oikos 104: 165–171.

Fortunel, C. et al. 2014. Environmental factors predict community functional composition in Amazonian forests. – J. Ecol. 102: 145–155.

Fox, J. and Weisberg, S. 2011. An {R} Companion to Applied Regression, Second Edition. Thousand Oaks CA: Sage. <http://socserv.socsci.mcmaster.ca/jfox/Books/Companion>.

Fowler, D. et al. 2014. Niche filtering rather than partitioning shapes the structure of temperate forest ant communities. – J. Anim. Ecol. 83: 943–952.

Fowler, H. G. et al. 1991. Ecologia nutricional de formigas. In: Panizzi, A.R. and Parra, J.R.P. (eds.). Ecologia nutricional de insetos. São Paulo, Editora Manole. p. 131–223.

Gerhold, P. et al. 2015. Phylogenetics patterns are not proxies of community assembly mechanisms (they are far better). – Funct. Ecol. 29: 600–614.

Gibb, H. and Cunningham, S. A. 2013. Restoration of trophic structure in an assemblage of omnivores, considering a revegetation chronosequence. – J. Appl. Ecol. 50: 449–458.

Gibb, H. and Parr, C. L. 2010. How does habitat complexity affect ant foraging success? A test of functional responses on three continents. – Oecologia 164: 1061–1073.

Gibb, H. and Parr, C. L. 2013. Does structural complexity determine the morphology of assemblages? An experimental test on three continents. – PLoS One 8: e64005.

Gibb, H. et al. 2015a. Climate mediates the effects of disturbance on ant assemblage structure. – Proc. R. Soc. Lond. B Biol. Sci. 282: 20150418.

Gibb, H. et al. 2015b. Does morphology predict trophic position and habitat use of ant species and assemblages? – Oecologia 177: 519–531.

Gotelli, N. J. and Ellison, A. M. 2002. Biogeography at a regional scale: determinants of ant species density in New England bogs and forests. – Ecology 83: 1604–1609.

Hijmans, R. J. et al. 2005. Very high resolution interpolated climate surfaces for global land areas. – Int. J. Climatol. 25: 1965–1978.

Inward, D. J. et al. 2011. Local and regional ecological morphology of dung beetle assemblage across four biogeography regions. – J. Biogeogr. 30: 1668–1682.

Jenkins, C. N. et al. 2011. Global diversity in ligh of climate change: the case of ants. – Divers. Distrib. 17: 652–662.

Kaspari, M. and Weiser, M. 1999. The size-grain hypothesis and interspecific scaling in ants. – Funct. Ecol. 13: 530–538.

Kraft, N. J. B. and Ackerly, D. D. 2010. Functional trait and phylogenetic tests of community assembly across spatial scales in an Amazonian forest. – Ecol. Monogr. 80: 401–422.

Kraft, N. J. B. et al. 2007. Trait evolution, community assembly, and the phylogenetic structure of ecological communities. – Am. Nat. 170: 271–283.

Kraft, N. J. B. et al. 2008. Functional traits and niche-based tree community assembly in an Amazonian forest. – Science 322: 580–582.

Lamanna, C. et al. 2014. Functional trait space and the latitudinal diversity gradient. – Proc. Natl. Acad. Sci. USA 111: 13745–13750.

Machac, A. et al. 2011. Elevational gradients in phylogenetic structure of ant communities reveal the interplay of biotic and abiotic constraints on diversity. – Ecography 34: 364–371.

McClain, C. R. et al. 2004. Morphological disparity as a biodiversity metric in lower bathyal and abyssal gastropod assemblages. – Evolution 58: 338–348.

McGill, B. J. et al. 2015. Fifteen forms of biodiversity trend in the Antropocene. – Trends Ecol. Evol. 30: 104–113.

Montaña, C. G. and Winemiller, K. O. 2010. Local-scale habitat influences morphological diversity of species assemblages of cichlid fishes in a tropical floodplain river. – Ecol. Freshw. Fish. 19: 216–227.

Montaña, C. G. et al. 2014. Intercontinental comparison of fish ecomorphology: null model tests of community assembly at the patch scale in rivers. – Ecol. Monogr. 84: 91–107.

Mouchet, M. A. et al. 2010. Functional diversity measures: an overview of their redundancy and their ability to discriminate community assembly rules. – Funct. Ecol. 24: 867–876.

Mouillot, D. et al. 2005. Niche overlap estimates based on quantitative functional traits: A new family of non-parametric indices. – Oecologia 145: 345–353.

Mouillot, D. et al. 2014. Functional over-redundancy and high functional vulnerability in global fish faunas on tropical reefs. – Proc. Natl. Acad. Sci. USA 111: 13757–13762.

Nipperess, D. A. and Beattie, A. J. 2004. Morphological dispersion of *Rhytidoponera* assemblages: the importance of spatial scale and null model. – Ecology 85: 2728–2736.

Petchey, O. L. and Gaston, K. J. 2002. Extinction and the loss of functional diversity. – Proc. R. Soc. Lond. B Biol. Sci. 269: 1721–1727.

Pie, M. R. and Traniello, J. F. A. 2007. Morphological evolution in a hyperdiverse clade: the ant genus *Pheidole*. – J. Zool. 271: 99–109.

Pinheiro J. et al. 2016. nlme: Linear and Nonlinear Mixed Effects Models. R package version 3.1–128, <http://CRAN.R-project.org/package=nlme>.

Raymond, A. et al. 2013. Functional diversity decreases with temperature in high elevation ant fauna. – Ecological Entomology 38: 364–373.

Ricklefs, R.E. 2012. Species richness and morphological diversity of passerine birds. – Proc. Natl. Acad. Sci. USA 109: 14482–14487.

Retana, J. et al. 2015. A multidimensional functional trait analysis of resource exploitation in European ants. – Ecology 96: 2781–2793.

Ricklefs, R. E. and Travis, J. 1980. A morphological approach to the study of avian community organization. – Auk 97: 321–338.

Sanders, N. J. et al. 2007. Assembly rules of ground-foraging ant assemblages are contingent on disturbance, habitat and spatial scale. – J. Biogeogr. 34: 1632–1641.

Sarty, M. et al. 2006. Habitat complexity facilities coexistence in a tropical ant community. – Oecologia 149: 465–473.

Schofield, S.F. et al. 2016. Morphological characteristics of ant assemblages (Hymenoptera: Formicidae) differ among contrasting biomes. – Myrmecol. News 23: 129–137.

Shmida, A. and Wilson, M. V. 1985. Biological determinants of species diversity. – J. Biogeogr. 12: 1–20.

Silva, R. R. and Brandão, C. R. F. 2010. Morphological patterns and community organization in leaf-litter ant assemblages. – Ecol. Monogr. 80: 107–124.

Silva, R. R. and Brandão, C. R. F. 2014. Ecosystem-wide morphological structure of leaf-litter ant communities along a tropical latitudinal gradient. – PLoS One 9: e93049.

Símová, I. et al. 2015. Shifts in trait means and variances in North American tree assemblages: species richness patterns are loosely related to the functional space. – Ecography 38: 649–658.

Spasojevic, M. J. and Suding, K. N. 2012. Inferring community assembly mechanisms from functional diversity patterns: the importance of multiple assembly mechanisms. – J. Ecol. 100: 652–661.

Stevens, R. D. et al. 2006. Latitudinal gradients in the phenetic diversity of New World bat communities. – Oikos 112: 41–50.

Stubbs, W. J. and Wilson, J. B. 2004. Evidence for limiting similarity in a sand dune community. – J. Ecol. 92: 557–567.

Stuble, K. L. et al. 2013. Tradeoffs, competition, and coexistence in eastern deciduous forest ant communities. – Oecologia 171: 981–992.

Swenson, N. G. 2014. Functional and Phylogenetic Ecology in R. Springer, New York. 212p.

Swenson, N. G. and Enquist, B. J. 2009. Opposing assembly mechanisms in a Neotropical dry forest: implications for phylogenetic and functional community ecology. – Ecology 90: 2161–2170.

Swenson, N. G. et al. 2012. The biogeography and filtering of woody plant functional diversity in North and South America trees. – Glob. Ecol. Biogeogr. 21: 798–808.

Swenson, N. G. and Weiser, M. D. 2014. On the packing and filling of functional space in eastern North American tree assemblages. – Ecography 37: 1056–1062.

Trisos, C. H. et al. 2014. Unraveling the interplay of community assembly processes acting on multiple niche axes across spatial scales. Am. Nat. 184: 593–608.

Venables, W. N. and Ripley, B. D. 2002. Modern Applied Statistics with S. Fourth Edition. Springer, New York.

Villéger, S. et al. 2008. New multidimensional functional diversity indices for a multifaceted framework in functional ecology. – Ecology 89: 2290–2301.

Webb, C. O. 2000. Exploring the phylogenetic structure of ecological communities: an example for rainforest trees. – Am. Nat. 156: 145–155.

Wiescher, P. T. et al. 2012. Assembling an ant community: species functional traits reflect environmental filtering. – Oecologia 169: 1063–1074.

Willis, S. C. et al. 2005. Habitat structural complexity and morphological diversity of fish assemblages in a Neotropical floodplain river. – Oecologia 142: 284–295.

Wilson, E. O. and Hölldobler, B. 2005. The rise of the ants: a phylogenetic and ecological explanation. – Proc. Natl. Acad. Sci. USA 102: 7411–7414.

Winemiller, K. O. 1991. Ecomorphological diversification in lowland freshwater fish assemblages from five biotic regions. – Ecol. Monogr. 61: 343–365.

Winemiller, K. O. et al. 2015. Functional traits, convergent evolution, and periodic tables of niches. – Ecol. Lett. 18: 737–751.

Yates, M. L. et al. 2014. Morphological traits: predictable responses to macrohabitats across a 300 km scale. – PeerJ 2: e271.

Yilmaz, A. et al. 2014. Eye structure, activity rhythms, and visually-driven behavior are tuned to visual niche in ants. – Front. Behav. Neurosci. 8: 1–9.

